# Development of a core genome multilocus sequence typing (cgMLST) scheme and life identification number (LIN) code classification system for *Staphylococcus aureus*

**DOI:** 10.1101/2025.03.29.646111

**Authors:** Nazreen F Hadjirin, Iman Yassine, James E Bray, Martin CJ Maiden, Keith A Jolley, Angela B Brueggemann

**Author notes:** Correspondence: Angela B Brueggemann.

## Abstract

*Staphylococcus aureus* infects both humans and animals, and antimicrobial resistance, including multidrug resistance, complicates treatment of *S. aureus* infections. Understanding *S. aureus* population structure and the distribution of genetic lineages is central to understanding the biology, epidemiology, and pathogenesis of this organism. This study exploited a large, publicly available dataset of nearly 27,000 *S. aureus* genomes to: i) develop a core genome multilocus sequence type (cgMLST) scheme; ii) stratify hierarchical clusters based on allelic similarity thresholds; and iii) define the clusters with a life identification number (LIN) code classification system. The cgMLST scheme characterised allelic variation at 1,716 core gene loci, and 13 classification thresholds were defined, which discriminated *S. aureus* variants across a range of genetic similarity thresholds. LIN code ‘lineages’ and clonal complexes defined by seven-locus multilocus sequence typing were highly concordant, but the LIN codes permitted a wider range of genetic discrimination among *S. aureus* genomes. This *S. aureus* cgMLST scheme and LIN code system is a high-resolution, stable genotyping tool that enables detailed genomic analyses of *S. aureus*.

## Introduction

*S. aureus* colonises healthy humans as well as healthy animals such as livestock, companion animals and wildlife, but is also associated with a wide range of diseases in both humans and animals. Globally, *S. aureus* caused an estimated 1.1 million human deaths in 2019 and over 100,000 of those deaths were attributable to methicillin-resistant *S. aureus* (MRSA) [1]. Antimicrobial resistance (AMR) among *S. aureus* is a major challenge worldwide, and the World Health Organisation has declared MRSA, vancomycin-intermediate, and vancomycin-resistant *S. aureus* to be a high AMR priority [2].

Bacterial whole genome sequence data are frequently used to investigate a wide range of questions related to population structure, epidemiology, infection, pathogenesis, and evolution. Genome sequencing is increasingly utilised in routine clinical microbiology, public health, and surveillance programmes [3]. The seven-locus multilocus sequence typing (MLST) scheme for *S. aureus* has become a standard genotyping methodology and the resulting sequence types (STs) are clustered into clonal complexes (CCs) for analyses [4,5]. The evolution and epidemiology of many of the major *S. aureus* CCs have been investigated in detail since the development of MLST and these findings have informed the overall understanding of this organism, especially in the context of the emergence and spread of MRSA [6-9].

However, seven-locus MLST has limited resolution, and genotyping tools that analyse more of the available sequence data can provide much more resolution, which has numerous applications. Such approaches include ribosomal MLST (rMLST), whereby genetic variation within the 53 bacterial ribosomal protein genes is characterised, which is especially useful for identifying bacterial species [10]. Core genome MLST (cgMLST) is used to characterise genetic variation among genes that are present in all or nearly all strains of a species, providing a much higher level of genetic resolution, since allelic variation among more than a thousand genes is typically used to define a core genome sequence type (cgST) for each genome. Gene-by-gene cgMLST schemes have been developed for a number of bacterial species [11-15]. A core genome typing scheme was developed for *S. aureus* in 2014, although new alleles for that scheme can only be submitted for assignment via a commercial platform, and a typing scheme called SaLTy was recently published that characterises just three core genes [16,17].

More recently, cgMLST schemes developed for *Klebsiella pneumoniae* and *Streptococcus pneumoniae* also included a new life identification number (LIN) barcoding classification system to cluster genomes using a range of similarity thresholds [13,14]. A major advantage of the new LIN code approach is that the resulting barcodes are fixed, and thus a stable hierarchical nomenclature is defined. Here, a publicly available cgMLST scheme and high-resolution LIN code system is described, which characterises *S. aureus* variants across multiple levels of genetic resolution, and the high-resolution clusters defined by cgSTs and LIN codes are compared to STs and CCs defined by seven-locus MLST.

## Materials and Methods

### Compilation of the PubMLST S. aureus genome database and study dataset

A total of 30,487 *S. aureus* genome assemblies were available in PubMLST at the time of this study. These had mostly been acquired in two ways (https://pubmlst.org/; accessed 15 May 2023). First, between 2012 and 2016, all available European Nucleotide Archive (ENA) high-throughput sequencing entries labelled as ‘ILLUMINA’ sequencing and organism as ‘Staphylococcus aureus’ (around 25,450 genomes) were assembled using Velvet/VelvetOptimiser and uploaded to the PubMLST *S. aureus* database (https://pubmlst.org/organisms/staphylococcus-aureus) [18-20]. Only minimal metadata were available for these genomes. Second, the remainder of the genomes and provenance data had been submitted by PubMLST database users between 2012 and 2023.

Genomes were chosen for inclusion in this study based on the following quality control criteria: i) total length, 2.6-3.0 Mb; ii) total number of contigs <300; iii) N50 value >25,000; iv) GC content, 32.0-33.1%; and v) an rMLST species designation of *S. aureus*. The rMLST species designations were performed by analysing each query genome relative to the rMLST genome library, as described previously [14]. In total, 26,677 genomes (330 complete genomes plus 26,347 draft genome assemblies) met the quality control criteria and were included (Supplementary Data 1). From this dataset, 5,000 genomes were randomly chosen as a development dataset, which included at least one representative of each of the 1,558 unique sequence types (STs; Supplementary Data 2 and 3). Randomisation was performed with the slice_sample in R version 4.2.3. CCs were defined using the global optimal eBURST algorithm implemented in PHYLOViZ v2.0.0 and CCs were named after the predicted ‘founder’ sequence type/s [5].

### *S. aureus* core genome loci and allele assignment

Previously, 2,265 core gene loci had been defined in PubMLST based on an early version of the *S. aureus* reference genome MRSA252 (NC_002952.1). These 2,265 loci were re-assessed for inclusion in a new cgMLST scheme, using the latest annotated version of the reference MRSA252 genome (NC_002952.2), to ensure that the cgMLST scheme only included non-paralogous core gene loci that were present in ≥99% of the 26,677 study genomes, and core gene loci without repeated sequence issues, in order to create a robust and reproducible cgMLST scheme.

Alleles were automatically assigned based on hits to the standard criteria of ≥97% sequence identity and 100% sequence alignment length to exemplar alleles defined for each locus such that all known alleles were within 97% identity of an exemplar of the same length. Core gene loci with missing allele assignments were manually curated and alleles were assigned wherever possible. Loci were excluded from the cgMLST scheme because of sequence quality issues (e.g. truncation, fragmentation, lacking a standard start or stop codon, variability in gene length, multiple allele assignments per assembly) and/or because alleles could not be assigned to at least 99% of the 26,677 study genomes. In total, after automated plus manual curation, 1,716 loci were included in the cgMLST scheme. Clusters of Orthologous Groups (COG) classifications were predicted using eggNOG-mapper v2 [21].

### Assignment of cgSTs and definition of LIN codes

All genomes that had 25 or fewer missing cgMLST alleles were assigned cgSTs using the BIGSdb automated profile definer. MSTclust (v0.21.200603ac; default parameters) was used to compute pairwise distances between cgMLST allelic profiles of the genomes in the development dataset (n = 5,000) and the full dataset (n = 26,677) [22]. MSTclust was also used to calculate the Silhouette (S_t_) score, a measure of cluster cohesiveness, from the 5,000 genomes using pairwise distances for the full range of values for every pairwise mismatch among the 1,716 loci.

The density distribution of pairwise allelic mismatches plus the Silhouette score were used to define a set of *S. aureus* classification thresholds using the development dataset of 5,000 genomes. A LIN code is a multi-position, integer-based barcode that uses the classification thresholds as ‘bins’ that correspond to cgMLST allelic mismatches. LIN codes for each distinct cgST were assigned within BIGSdb as described previously [13,14]. The concordance between cgST clusters versus other *S. aureus* molecular typing methods (CC, ST, ribosomal sequence type [rST], SaLTy) was tested by calculating the adjusted Rand Index for each classification threshold using the development dataset of 5,000 genomes. The analysis was then repeated using the full dataset to test the concordance between cgMLST and CC, ST, rST [23].

### Phylogenetic analyses

A subset of 1,558 genomes, representing one example of each unique ST, was used to construct a maximum likelihood phylogenetic tree with the 1,716 concatenated core gene sequences, to test the congruence of the phylogenetic tree topology relative to each of the defined LIN code thresholds. Allele sequences for the 1,716 cgMLST loci in each genome were retrieved from the BIGSdb sequence definition database, aligned using MAFFT (v7.515; missing alleles were treated as gaps) and concatenated [24]. snp-sites (v2.5.1) was used to calculate the number of variant sites in the sequence alignments [25]. Maximum likelihood phylogenetic trees were reconstructed with IQ-TREE (v2.0.3), and the built-in model testing (-m MFP) with 1000 ultrafast bootstraps (-bb 1000) was used to determine the best phylogenetic model [26]. Trees were midpoint rooted and visualised in ggtree or iTOL [27,28]. Minimum spanning trees were constructed using GrapeTree and the allelic profiles of the 1,716 core genes [29].

## Results

### Description of the S. aureus genome dataset

The country of origin and year of isolation were available in PubMLST for approximately half of the genomes (54.4% and 48.4% respectively), indicating that *S. aureus* from 72 countries and 6 continents were represented, and they were recovered between 1930 and 2023 (Supplementary Data 4). Around two-thirds of the 14,510 *S. aureus* with a known country of origin were either from the USA (45.8%, n = 6,645) or the United Kingdom (16.7%, n = 2,420). The original source of the *S. aureus* isolate was available for only 11.9% (n = 3,172) of the genome dataset, of which 1,886 were from humans and 1,120 were from animals (Supplementary Data 4). In total, 5,440 unique rSTs and 1,558 unique STs were represented, and the 1,558 STs clustered into 90 CCs consisting of 2 to 6,405 members per CC. Collectively, 20 CCs represented 95.6% (n = 25,510) of all genomes in the dataset (Table 1; Supplementary Data 4).

**Table 1.** Twenty clonal complexes (CCs) represented more than 95% of the genomes in the S aureus dataset.

| CC | Genomes (n) | % of dataset |
| --- | --- | --- |
| 5 | 6,405 | 24.0 |
| 8 | 5,547 | 20.8 |
| 22 | 5,107 | 19.1 |
| 398 | 1,700 | 6.4 |
| 30 | 1,514 | 5.7 |
| 1 | 903 | 3.4 |
| 45 | 683 | 2.6 |
| 130 | 549 | 2.1 |
| 15 | 530 | 2.0 |
| 93 | 469 | 1.8 |
| 97 | 384 | 1.4 |
| 133 | 313 | 1.2 |
| 9 | 277 | 1.0 |
| 59 | 243 | 0.9 |
| 188 | 211 | 0.8 |
| 425 | 167 | 0.6 |
| 88 | 143 | 0.5 |
| 121 | 134 | 0.5 |
| 25 | 126 | 0.5 |
| 72 | 105 | 0.4 |
| **Total<sup>a</sup>** | **25,510** | **95.6** |

### The cgMLST genotyping scheme

The 1,716 loci included in the cgMLST scheme were distributed across the *S. aureus* genome and 72.7% (n = 1,248) were predicted to encode proteins with COG classified cellular pathways (Supplementary Figure 1; Supplementary Data 5). Overall, 98.8% (n = 26,347) of the 26,677 *S. aureus* genomes had 25 or fewer missing cgST alleles and thus were assigned a cgST. There were 23,136 unique cgSTs (Supplementary Data 6).

Among genomes with 25 or fewer missing cgST alleles, the mean number of missing alleles was comparable across CCs, suggesting little phylogenetic bias (Supplementary Figure 3). Among the 330 genomes that had more than 25 missing alleles the median was 37 (range, 26-138) missing alleles, but these were not assigned a cgST and LIN code to prevent isolates appearing to be artefactually closer to each other than they really are, due to missing loci being ignored in pairwise comparisons.

### Determination of S. aureus classification thresholds

The development dataset of 5,000 genomes was used to define thresholds of genetic similarity to delineate the *S. aureus* population structure (Figure 1). A multimodal distribution of pairwise allelic mismatches among the 1,716 cgMLST loci was observed, which included three major peaks centred at 94%, 89% and 80%, and minor peaks around 30%, 13% and 4% allelic mismatches.

**Figure 1.**
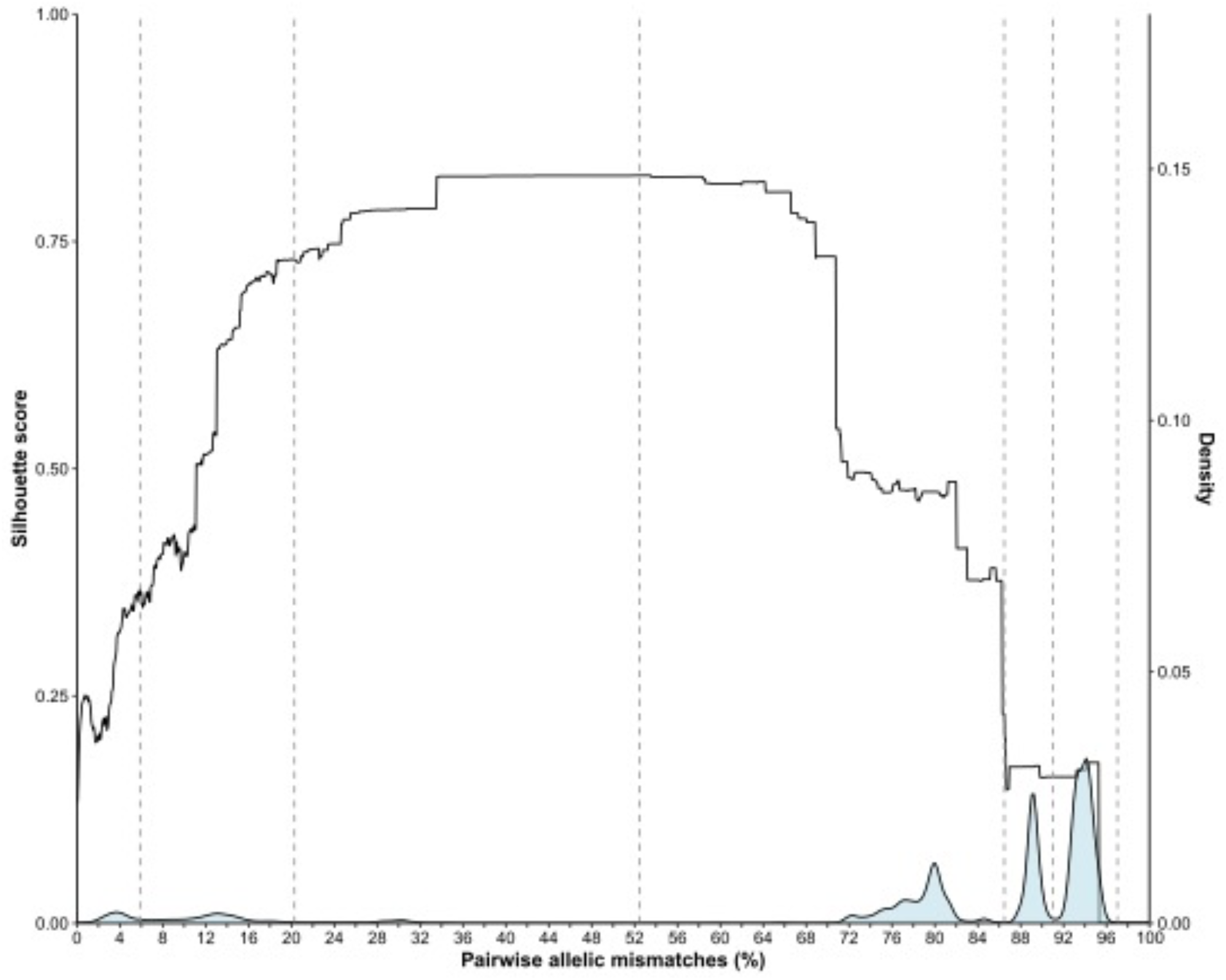
Determination of cgMLST classification thresholds to define the population structure of *S. aureus*. The development dataset of 5,000 genomes, which included at least one example of each of 1,558 unique seven-locus STs, was used in these analyses. Vertical lines mark the first six LIN code classification thresholds. Note that the final seven classification thresholds on the left are positioned very closely together and thus not illustrated here. cgMLST, core genome multilocus sequence typing; STs, sequence types.

A total of 13 thresholds were chosen to provide a range of discrimination levels (Figure 1). The first threshold was defined at 1,665 pairwise allelic mismatches (i.e. 97.0% genetic dissimilarity): this was the maximum number of core gene allelic differences among all 5,000 genomes. The second threshold was set at 1,562 (91.0%) allelic mismatches to split two major peaks in the distribution. A third threshold was set at 1,484 (86.5%) allelic mismatches to flank the second major peak and coincide with the increasing Silhouette score. A ‘lineage’ threshold was set at the highest Silhouette score (S_t_=0.82) and defined genomes with 900 (52.4%) pairwise allelic mismatches. Near the minor peaks, a fifth threshold was set at 348 (20.3%) allelic mismatches, and a sixth threshold was defined at 101 (5.9%) allelic mismatches. Finally, high-level discrimination thresholds were set at 27 (1.6%), 14 (0.8%), 7 (0.4%), 4 (0.2%), 2 (0.1%), 1 (0.06%) and 0 (0%) pairwise allelic mismatches.

### Comparison of the cgST-LIN code scheme with other S. aureus genotyping methods

Expanding the analyses to the dataset of 26,347 genomes with a cgST, the distributions of pairwise allelic differences between genomes belonging to a matching rST, ST or CC group were assessed. The majority of genomes that had matching STs or CCs had fewer than 30% allelic mismatches among the cgST loci, although there was a small group of genomes within CC1, CC5, and CC188 that had matching CCs but around 75-80% allelic mismatches at core gene loci (Figure 2A).

**Figure 2.**
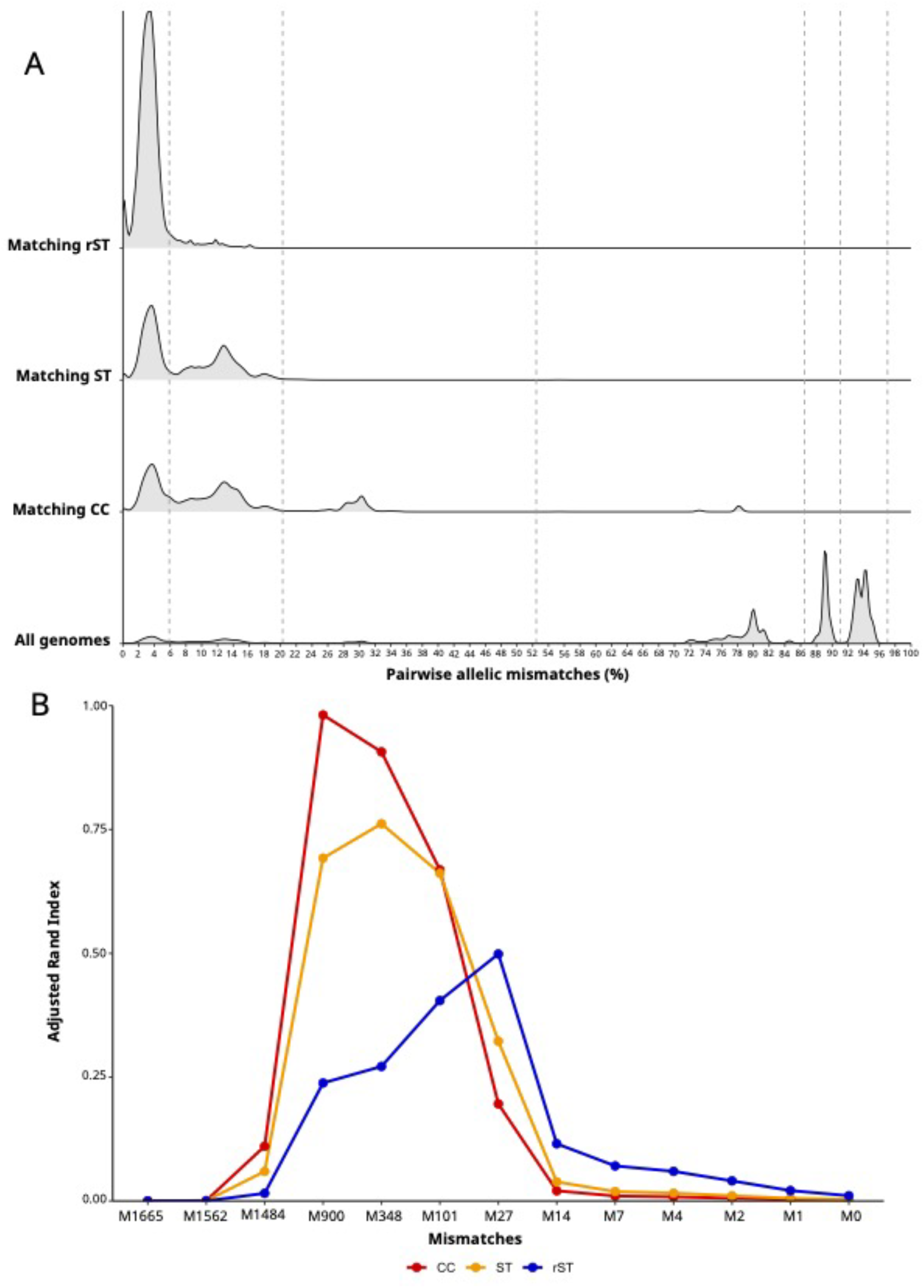
Characterisation of *S. aureus* genomes by cgMLST and comparison to other genotyping methods. The results of the analysis of 26,347 *S. aureus* genomes with an assigned cgST are illustrated here. Panel A depicts the density distributions of pairwise allelic mismatches between genomes belonging to a matching rST, ST or CC taxonomic group. Panel B plots the concordance between clusters of *S aureus* at each of the 13 LIN code thresholds (marked here by the number of pairwise mismatches) and the corresponding rST, ST and CC designations. The fourth threshold at 900 pairwise mismatches was designed a LIN code ‘lineage’. Note: cgMLST, core genome multilocus sequence typing; ST, sequence type; cgST, core genome ST; rST, ribosomal ST; CC, clonal complex; M, mismatches.

The concordance between clusters of *S. aureus* at each of the LIN code thresholds and the corresponding rST, ST and CC designations was calculated using the adjusted Rand Index (ARI). There was nearly perfect concordance (ARI = 0.98) between the LIN code ‘lineage’ (900 pairwise mismatches) and CC, high concordance (ARI = 0.76) between LIN code threshold 5 (348 pairwise mismatches) and ST, and moderate concordance (ARI = 0.50) between LIN code threshold 7 (27 pairwise mismatches) and rST (Figure 2B). (A similar analysis that included SaLTy was run on the development dataset of 5,000 genomes; Supplementary Figure 3).

### Increased resolution of S. aureus population structure using cgMLST and LIN codes

A maximum likelihood phylogenetic tree was constructed using concatenated nucleotide sequence alignments of the 1,716 cgMLST loci of one randomly chosen genome representative for each of the 1,558 unique STs, and the tree was annotated with the 20 most common CCs in this dataset and their corresponding LIN code lineages. This analysis clearly demonstrated concordance between CC and LIN code lineage (Figure 3; Supplementary Figure 4).

**Figure 3.**
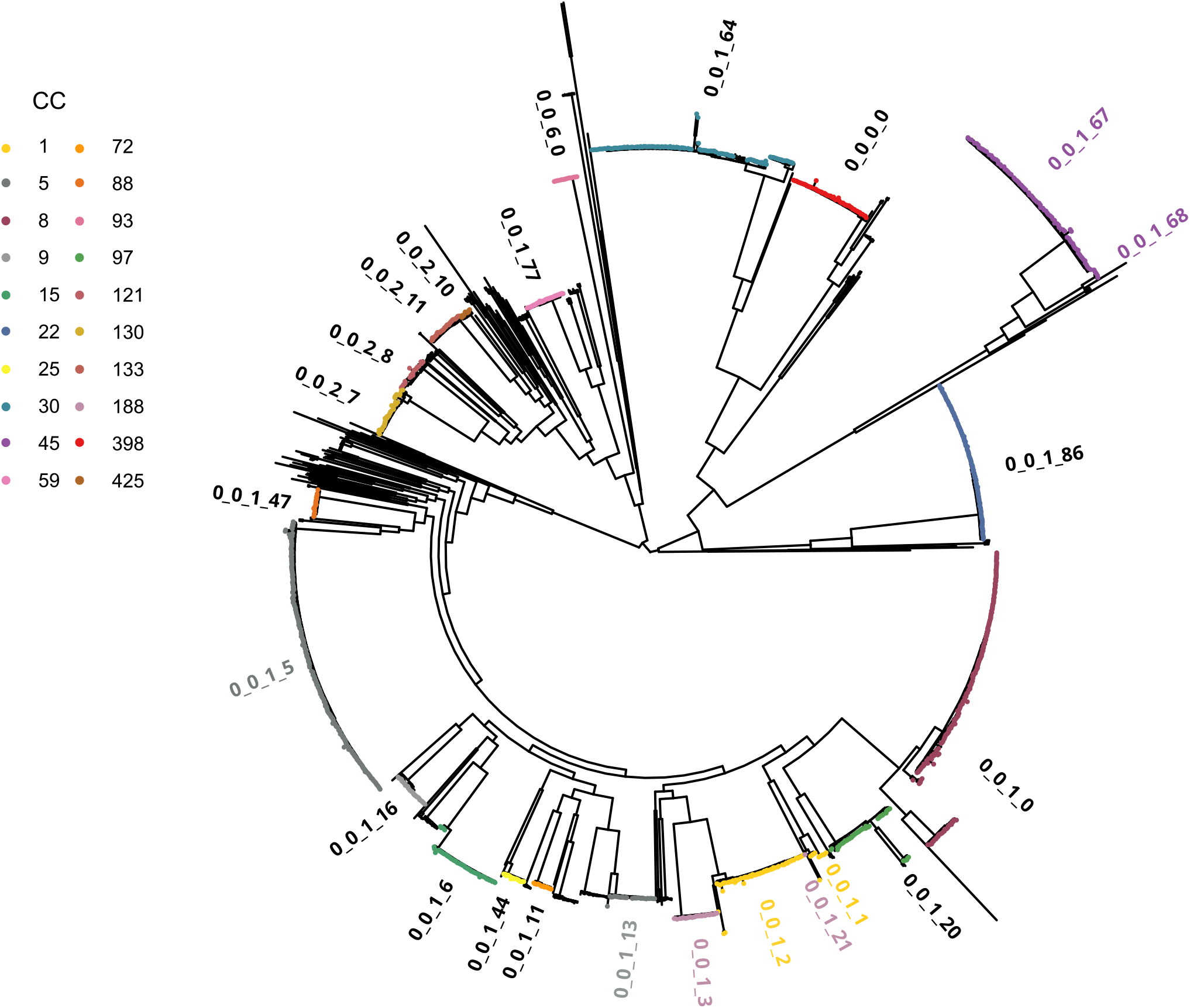
Maximum-likelihood phylogeny showing the population structure of *S. aureus*. The tree was constructed with a concatenated alignment of 1,716 core genome loci from one randomly selected genome representative for each of the 1,558 unique seven-locus sequence types. IQ-TREE was used with the GTR+G model of nucleotide substitution and 1,000 bootstrap replicates. The tree is rooted at the midpoint. The 20 most common CCs in the dataset are coloured and annotated with the corresponding LIN code classification threshold 4 ‘lineage’ designations. Note that CCs 1, 5, 45 and 188 were split into two LIN code lineages each (see main text and Figures 4 and 5).

Among the 26,347 genomes with an assigned cgST and LIN code, there were 18,455 unique LIN codes. There were 10 LIN code 3 thresholds (8 of which represented more than one genome; range, 3 to 22,727), 119 LIN code lineages (74 of which represented more than one genome; range, 2 to 6,298), and 167 LIN code 5 thresholds (96 of which represented more than one genome; range, 2 to 6,298) (Supplementary Data 6).

Twenty-two LIN code lineages comprised 96.4% (n = 25,411) of the 26,347 *S. aureus* genomes with LIN codes. These 22 LIN code lineages also represented the 20 most common CCs in the dataset; however, CC5 *S. aureus* were represented by two LIN code lineages 0_0_1_5 (n = 6,298 genomes) and 0_0_1_13 (n = 125 genomes), and CC1 *S. aureus* were roughly evenly split into two LIN code lineages 0_0_1_2 (n = 511 genomes) and 0_0_1_1 (n = 398 genomes), respectively (Table 2). There were two additional CCs that were also split into two LIN code lineages, but only one of each LIN code was represented in >95% of the full dataset: CC45 (0_0_1_67, n = 637 genomes; 0_0_1_68, n = 46 genomes); and CC188 (0_0_1_3, n = 201 genomes; 0_0_1_21, n = 1 genome; Table 2, Supplementary Data 6).

**Table 2.** Twenty-two LIN code lineages represented more than 95% of the genomes in the *S aureus* dataset.

| **LIN code lineage** | **Genomes in lineage (n)** | **% of dataset<sup>a</sup>** | **Major CC** | **Number (%) of genomes in lineage that are also in the major CC** |
| --- | --- | --- | --- | --- |
| 0_0_1_5 | 6,298 | 23.9 | CC5 | 6,231 (98.9) |
| 0_0_1_0 | 5,539 | 21.0 | CC8 | 5,516 (99.6) |
| 0_0_1_86 | 5,083 | 19.3 | CC22 | 5,080 (99.9) |
| 0_0_0_0 | 1,680 | 6.4 | CC398 | 1,679 (99.9) |
| 0_0_1_64 | 1,557 | 5.9 | CC30 | 1,504 (96.6) |
| 0_0_1_67 | 637 | 2.4 | CC45 | 633 (99.4) |
| 0_0_2_7 | 557 | 2.1 | CC130 | 546 (98.0) |
| 0_0_1_6 | 518 | 2.0 | CC15 | 518 (100) |
| 0_0_1_2 | 511 | 1.9 | CC1 | 501 (98.0) |
| 0_0_6_0 | 467 | 1.8 | CC93 | 467 (100) |
| 0_0_1_1 | 398 | 1.5 | CC1 | 397 (99.7) |
| 0_0_1_20 | 337 | 1.3 | CC97 | 332 (98.5) |
| 0_0_2_11 | 312 | 1.2 | CC133 | 312 (100) |
| 0_0_1_16 | 270 | 1.0 | CC9 | 270 (100) |
| 0_0_1_77 | 240 | 0.9 | CC59 | 238 (99.2) |
| 0_0_1_3 | 201 | 0.8 | CC188 | 200 (99.5) |
| 0_0_2_10 | 164 | 0.6 | CC425 | 164 (100) |
| 0_0_2_8 | 149 | 0.6 | CC121 | 131 (87.9) |
| 0_0_1_47 | 145 | 0.6 | CC88 | 142 (97.9) |
| 0_0_1_13 | 125 | 0.5 | CC5 | 122 (97.6) |
| 0_0_1_44 | 121 | 0.5 | CC25 | 120 (99.2) |
| 0_0_1_11 | 102 | 0.4 | CC72 | 102 (100) |
| **Total<sup>a</sup>** | **25,411** | **96.4** | **--** | **--** |

a. Note that there were 26,347 genomes with an assigned LIN code in the study dataset.

The relationships among the CC5 and CC1 pairs of LIN code lineages were visualised with GrapeTree, which used the allelic profiles of the 1,716 core gene loci as input data (Figures 4 and 5). Within each pair of LIN code lineages, the two clusters of *S. aureus* were distinct (despite being in the same CC), and the increased genetic resolution provided by LIN codes was visualised by annotating the tree at three higher LIN code classification thresholds.

**Figure 4.**
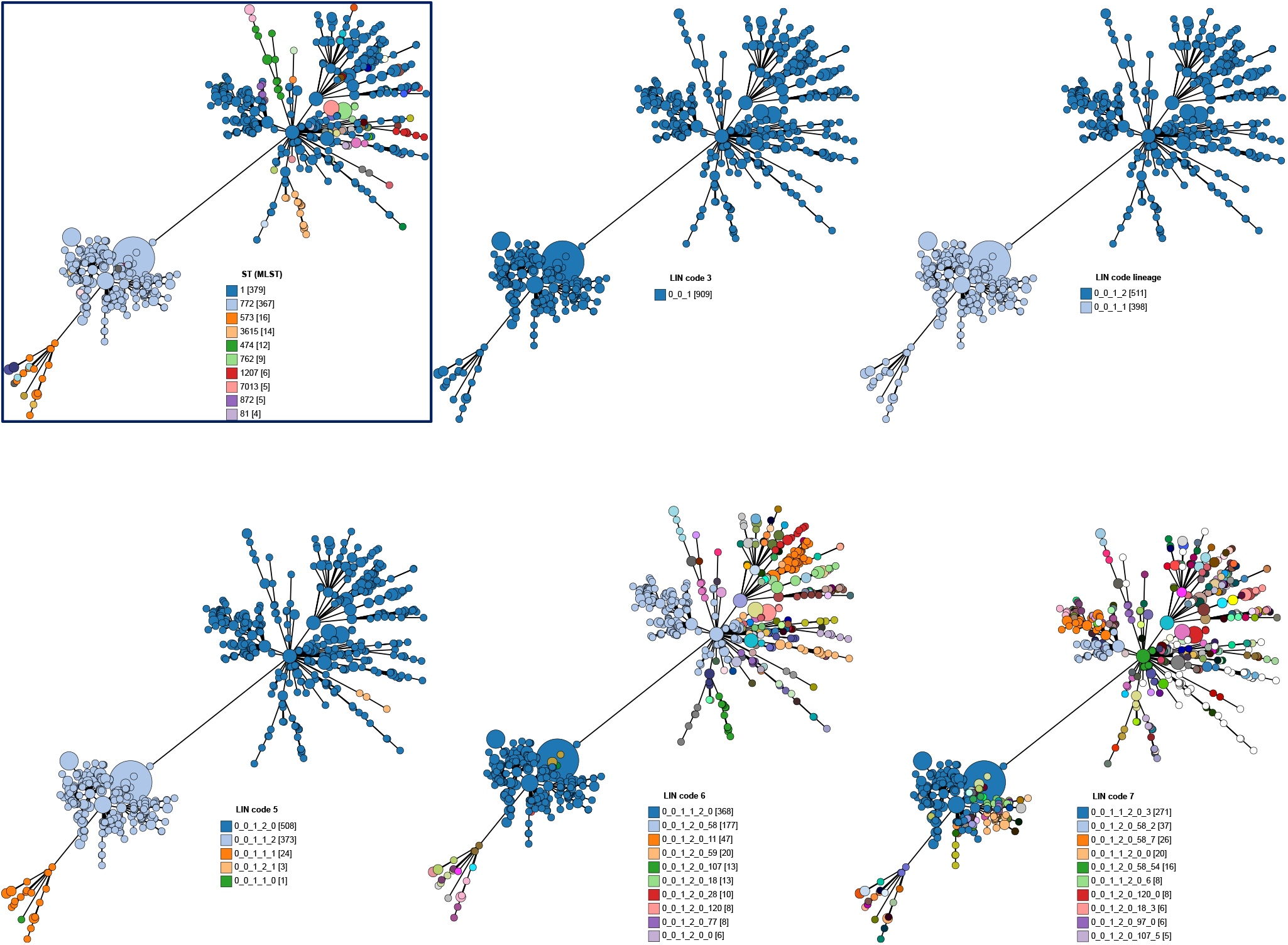
*S. aureus* members of clonal complex 1 were split into two LIN code lineages. Relationships between the two LIN code lineages 0_0_1_2 (n = 511) and 0_0_1_1 (n = 398) were visualised with GrapeTree, which plots allelic profiles of the cgMLST loci. The same minimum spanning tree is shown in each panel, coloured by ST or LIN code. The *S. aureus* in the tree in the boxed panel were coloured by seven-locus sequence type (ST), and the remaining panels were coloured by increasing LIN code classification thresholds. In the legends, the numbers in square brackets indicate the number of *S. aureus* with that ST or LIN code. For illustrative purposes, the legend only shows the 10 most common LIN codes annotated in the LIN code 6 and LIN code 7 trees.

**Figure 5.**
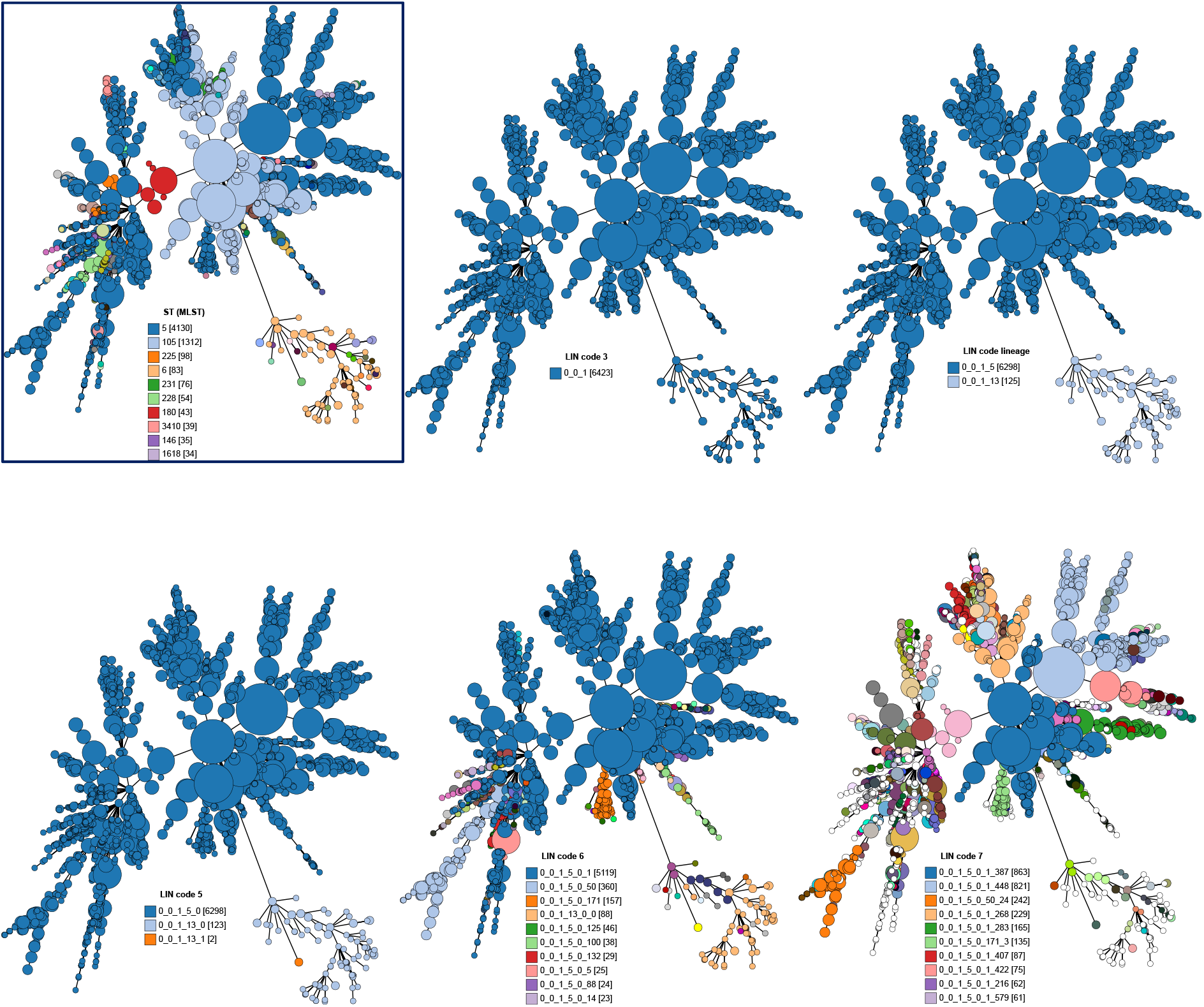
*S. aureus* members of clonal complex 5 were split into two LIN code lineages. Relationships between the two LIN code lineages 0_0_1_5 (n = 6,298) and 0_0_1_13 (n = 125) were visualised with GrapeTree, which plots allelic profiles of the cgMLST loci. The same minimum spanning tree is shown in each panel, coloured by ST or LIN code. The *S. aureus* in the tree in the boxed panel were coloured by seven-locus sequence type (ST), and the remaining panels were coloured by increasing LIN code classification thresholds. In the legends, the numbers in square brackets indicate the number of *S. aureus* with that ST or LIN code. For illustrative purposes, the legend only shows the 10 most common LIN codes annotated in the LIN code 6 and LIN code 7 trees.

## Discussion

Robust genotyping tools for bacterial pathogens are essential for microbiological research and public health surveillance, and a well-defined, internationally recognised nomenclature is critical for sharing and communicating the genotyping results within the scientific community [30]. The seven-locus MLST scheme has proven to be a robust and widely used typing tool for defining STs and CCs of *S. aureus*, in particular because S. aureus has relatively low rates of recombination and the CCs tend to be stable [7,31]; however, whole genome sequencing enables the characterisation of bacterial variants at much greater resolution than seven-locus MLST. The publicly available cgMLST scheme and LIN code system described here provides a stable and precise typing scheme for *S. aureus*.

One of the challenges of proposing a new classification system for bacterial species with existing nomenclatures is achieving consistency with currently accepted schema. The *S. aureus* cgMLST scheme and LIN code system proposed here mitigates this by having backwards compatibility with the major CCs, and the PubMLST website and BIGSdb platform provide the relevant nomenclature. A 13 hierarchical LIN code classification threshold system has been established that delineates the *S. aureus* population structure, enabling the user to choose the level of resolution they require. The designated LIN code ‘lineage’ is highly concordant with CC, and LIN code threshold 5 is largely concordant with ST, providing consistency across the available genotyping methods.

The *S. aureus* genome database was initially established by downloading and assembling sequencing data from the ENA, and relatively limited provenance data and phenotype data were available for each genome sequence. This has limited effects on the development of a robust genotyping scheme but does preclude a more detailed investigation into the epidemiology and evolution of defined genetic clusters. More recent submissions include more detailed and complete provenance data, and we encourage future submitters of genome data to PubMLST to provide as much provenance data as possible, for the benefit of the wider community.

In conclusion, a high-resolution, open access cgMLST scheme and LIN code system is proposed, which enables the detailed analyses of genetic lineages within the *S. aureus* population. The varying levels of resolution provided by these new genotyping tools will be advantageous for *S. aureus* outbreak investigations, epidemiological surveillance, and addressing research questions related to *S. aureus* evolution, pathogenesis, and antimicrobial resistance.

## Data availability

All genome sequences and corresponding metadata are available from PubMLST (https://pubmlst.org/organisms/staphylococcus-aureus). Genome accession numbers are available within each isolate record in PubMLST and in Supplementary Data 1.

## Code availability

The relevant code for data analyses can be found here: https://github.com/brueggemann-lab/pgl_cgmlst_2024

## Authors and contributors

Conceptualisation: NFH, KAJ, ABB. Data curation: NFH, JEB, KAJ. Genome assembly: JEB. Data analyses: NFH, IY, JEB, KAJ, ABB. Data visualisation: NFH, IY, KAJ, ABB. PubMLST platform and software development: JEB, MCJM, KAJ, ABB. PubMLST funding and infrastructure: MCJM, KAJ, ABB. Writing of first draft: NFH, ABB. All authors contributed to the final version of the manuscript.

## Acknowledgements

The authors thank Duncan Berger for helpful discussions about the analyses.

## References

1. Antimicrobial Resistance Collaborators. Global burden of bacterial antimicrobial resistance in 2019: a systematic analysis. Lancet. 2022 Feb 12;399(10325):629–655. doi: 10.1016/S0140-6736(21)02724-0.

2. Estimating the impact of vaccines in reducing antimicrobial resistance and antibiotic use: technical report. Geneva: World Health Organization; 2024. https://iris.who.int/

3. Besser J, Carleton HA, Gerner-Smidt P, Lindsey RL, Trees E. Next-generation sequencing technologies and their application to the study and control of bacterial infections. Clin Microbiol Infect. 2018 Apr;24(4):335–341. doi: 10.1016/j.cmi.2017.10.013.

4. Enright MC, Day NP, Davies CE, Peacock SJ, Spratt BG. Multilocus sequence typing for characterization of methicillin-resistant and methicillin-susceptible clones of Staphylococcus aureus. J Clin Microbiol. 2000 Mar;38(3):1008–15. doi: 10.1128/JCM.38.3.1008-1015.2000.

5. Nascimento M, Sousa A, Ramirez M, Francisco AP, Carriço JA, Vaz C. PHYLOViZ 2.0: providing scalable data integration and visualization for multiple phylogenetic inference methods. Bioinformatics. 2017 Jan 1;33(1):128–129. doi: 10.1093/bioinformatics/btw582.

6. Richardson EJ, Bacigalupe R, Harrison EM, Weinert LA, Lycett S, Vrieling M, Robb K, Hoskisson PA, Holden MTG, Feil EJ, Paterson GK, Tong SYC, Shittu A, van Wamel W, Aanensen DM, Parkhill J, Peacock SJ, Corander J, Holmes M, Fitzgerald JR. Gene exchange drives the ecological success of a multi-host bacterial pathogen. Nat Ecol Evol. 2018 Sep;2(9):1468–1478. doi: 10.1038/s41559-018-0617-0.

7. Feil EJ, Cooper JE, Grundmann H, Robinson DA, Enright MC, Berendt T, Peacock SJ, Smith JM, Murphy M, Spratt BG, Moore CE, Day NP. How clonal is Staphylococcus aureus? J Bacteriol. 2003 Jun;185(11):3307–16. doi: 10.1128/JB.185.11.3307-3316.2003.

8. Holden MT, Hsu LY, Kurt K, Weinert LA, Mather AE, Harris SR, Strommenger B, Layer F, Witte W, de Lencastre H, Skov R, Westh H, Zemlicková H, Coombs G, Kearns AM, Hill RL, Edgeworth J, Gould I, Gant V, Cooke J, Edwards GF, McAdam PR, Templeton KE, McCann A, Zhou Z, Castillo-Ramírez S, Feil EJ, Hudson LO, Enright MC, Balloux F, Aanensen DM, Spratt BG, Fitzgerald JR, Parkhill J, Achtman M, Bentley SD, Nübel U. A genomic portrait of the emergence, evolution, and global spread of a methicillin-resistant Staphylococcus aureus pandemic. Genome Res. 2013 Apr;23(4):653–64. doi: 10.1101/gr.147710.112.

9. McAdam PR, Templeton KE, Edwards GF, Holden MT, Feil EJ, Aanensen DM, Bargawi HJ, Spratt BG, Bentley SD, Parkhill J, Enright MC, Holmes A, Girvan EK, Godfrey PA, Feldgarden M, Kearns AM, Rambaut A, Robinson DA, Fitzgerald JR. Molecular tracing of the emergence, adaptation, and transmission of hospital-associated methicillin-resistant Staphylococcus aureus. Proc Natl Acad Sci U S A. 2012 Jun 5;109(23):9107–12. doi: 10.1073/pnas.1202869109.

10. Jolley KA, Bliss CM, Bennett JS, Bratcher HB, Brehony C, Colles FM, Wimalarathna H, Harrison OB, Sheppard SK, Cody AJ, Maiden MCJ. Ribosomal multilocus sequence typing: universal characterization of bacteria from domain to strain. Microbiology (Reading). 2012 Apr;158(Pt 4):1005–1015. doi: 10.1099/mic.0.055459-0.

11. Liang KYH, Orata FD, Islam MT, Nasreen T, Alam M, Tarr CL, Boucher YF. A Vibrio cholerae core genome multilocus sequence typing scheme to facilitate the epidemiological study of cholera. J Bacteriol. 2020 Nov 19;202(24):e00086–20. doi: 10.1128/JB.00086-20.

12. Yassine I, Lefèvre S, Hansen EE, Ruckly C, Carle I, Lejay-Collin M, Fabre L, Rafei R, Clermont D, de la Gandara MP, Dabboussi F, Thomson NR, Weill FX. Population structure analysis and laboratory monitoring of Shigella by core-genome multilocus sequence typing. Nat Commun. 2022 Jan 27;13(1):551. doi: 10.1038/s41467-022-28121-1.

13. Hennart M, Guglielmini J, Bridel S, Maiden MCJ, Jolley KA, Criscuolo A, Brisse S. A dual barcoding approach to bacterial strain nomenclature: genomic taxonomy of Klebsiella pneumoniae strains. Mol Biol Evol. 2022 Jul 2;39(7):msac135. doi: 10.1093/molbev/msac135. PMID: 35700230; PMCID: PMC9254007.

14. Jansen van Rensburg MJ, Berger DJ, Yassine I, Shaw D, Fohrmann A, Bray JE, Jolley KA, Maiden MCJ, Brueggemann AB. Development of the Pneumococcal Genome Library, a core genome multilocus sequence typing scheme, and a taxonomic life identification number barcoding system to investigate and define pneumococcal population structure. Microb Genom. 2024 Aug;10(8):001280. doi: 10.1099/mgen.0.001280.

15. Krisna MA, Jolley KA, Monteith W, Boubour A, Hamers RL, Brueggemann AB, Harrison OB, Maiden MCJ. Development and implementation of a core genome multilocus sequence typing scheme for Haemophilus influenzae. Microb Genom. 2024 Aug;10(8):001281. doi: 10.1099/mgen.0.001281.

16. Leopold SR, Goering RV, Witten A, Harmsen D, Mellmann A. Bacterial whole-genome sequencing revisited: portable, scalable, and standardized analysis for typing and detection of virulence and antibiotic resistance genes. J Clin Microbiol. 2014 Jul;52(7):2365–70. doi: 10.1128/JCM.00262-14.

17. Cheney L, Payne M, Kaur S, Lan R. SaLTy: a novel Staphylococcus aureus Lineage Typer. Microb Genom. 2024 May;10(5):001250. doi: 10.1099/mgen.0.001250.

18. Zerbino DR, Birney E. Velvet: algorithms for de novo short read assembly using de Bruijn graphs. Genome Res. 2008 May;18(5):821–9. doi: 10.1101/gr.074492.107.

19. https://github.com/tseemann/VelvetOptimiser

20. Jolley KA, Bray JE, Maiden MCJ. Open-access bacterial population genomics: BIGSdb software, the PubMLST.org website and their applications. Wellcome Open Res. 2018 Sep 24;3:124. doi: 10.12688/wellcomeopenres.14826.1.

21. Cantalapiedra CP, Hernández-Plaza A, Letunic I, Bork P, Huerta-Cepas J. eggNOG-mapper v2: Functional Annotation, Orthology Assignments, and Domain Prediction at the Metagenomic Scale. Mol Biol Evol. 2021 Dec 9;38(12):5825–5829. doi: 10.1093/molbev/msab293.

22. https://gitlab.pasteur.fr/GIPhy/MSTclust

23. Rand, W.M. Objective Criteria for the Evaluation of Clustering Methods. J Am Stat Assoc 1971; 66, 846–850.

24. Katoh K, Misawa K, Kuma K, Miyata T. MAFFT: a novel method for rapid multiple sequence alignment based on fast Fourier transform. Nucleic Acids Res. 2002 Jul 15;30(14):3059–66. doi: 10.1093/nar/gkf436.

25. https://sanger-pathogens.github.io/snp-sites/

26. Nguyen LT, Schmidt HA, von Haeseler A, Minh BQ. IQ-TREE: a fast and effective stochastic algorithm for estimating maximum-likelihood phylogenies. Mol Biol Evol. 2015 Jan;32(1):268–74. doi: 10.1093/molbev/msu300.

27. Yu, G., Smith, D.K., Zhu, H., Guan, Y. and Lam, T.T.-Y. (2017), GGTREE: an R package for visualization and annotation of phylogenetic trees with their covariates and other associated data. Methods Ecol Evol, 8: 28–36. 10.1111/2041-210X.12628

28. Letunic I, Bork P. Interactive Tree of Life (iTOL) v6: recent updates to the phylogenetic tree display and annotation tool. Nucleic Acids Res. 2024 Jul 5;52(W1):W78–W82. doi: 10.1093/nar/gkae268.

29. Zhou Z, Alikhan NF, Sergeant MJ, Luhmann N, Vaz C, Francisco AP, Carriço JA, Achtman M. GrapeTree: visualization of core genomic relationships among 100,000 bacterial pathogens. Genome Res. 2018 Sep;28(9):1395–1404. doi: 10.1101/gr.232397.117.

30. Maiden MC, Jansen van Rensburg MJ, Bray JE, Earle SG, Ford SA, Jolley KA, McCarthy ND. MLST revisited: the gene-by-gene approach to bacterial genomics. Nat Rev Microbiol. 2013 Oct;11(10):728–36. doi: 10.1038/nrmicro3093.

31. Everitt RG, Didelot X, Batty EM, Miller RR, Knox K, Young BC, Bowden R, Auton A, Votintseva A, Larner-Svensson H, Charlesworth J, Golubchik T, Ip CL, Godwin H, Fung R, Peto TE, Walker AS, Crook DW, Wilson DJ. Mobile elements drive recombination hotspots in the core genome of Staphylococcus aureus. Nat Commun. 2014 May 23;5:3956. doi: 10.1038/ncomms4956.

