## Supplementary Table for "Development of a core genome multilocus sequence typing (cgMLST) scheme and life identification number (LIN) code classification system for *Staphylococcus aureus*"

**Supplementary Table 1.** List of all clonal complexes and singletons^a^ in the full *S. aureus* study dataset.

| CC | Genomes (n) | % of dataset |
| --- | --- | --- |
| 5 | 6405 | 24.0 |
| 8 | 5547 | 20.8 |
| 22 | 5107 | 19.1 |
| 398 | 1700 | 6.4 |
| 30 | 1514 | 5.7 |
| 1 | 903 | 3.4 |
| 45 | 683 | 2.6 |
| 130 | 549 | 2.1 |
| 15 | 530 | 2.0 |
| 93 | 469 | 1.8 |
| 97 | 384 | 1.4 |
| 133 | 313 | 1.2 |
| 9 | 277 | 1.0 |
| 59 | 243 | 0.9 |
| 188 | 211 | 0.8 |
| 425 | 167 | 0.6 |
| 88 | 143 | 0.5 |
| 121 | 134 | 0.5 |
| 25 | 126 | 0.5 |
| 72 | 105 | 0.4 |
| 12 | 97 | 0.4 |
| 7 | 91 | 0.3 |
| 151/705 | 90 | 0.3 |
| 228/111 | 66 | 0.2 |
| 49 | 59 | 0.2 |
| 20 | 57 | 0.2 |
| 39 | 48 | 0.2 |
| 101 | 42 | 0.2 |
| 1943 | 39 | 0.1 |
| 3881/3894 | 32 | 0.1 |
| 522 | 32 | 0.1 |
| 80 | 31 | 0.1 |
| 672 | 31 | 0.1 |
| 152 | 30 | 0.1 |
| 182 | 29 | 0.1 |
| 291 | 28 | 0.1 |
| 50 | 22 | 0.1 |
| 123 | 16 | 0.1 |
| 126 | 13 | 0.0 |
| 692 | 12 | 0.0 |
| 599/2179/2508 | 11 | 0.0 |
| 5993 | 9 | 0.0 |
| 1797 | 9 | 0.0 |
| 479/520 | 8 | 0.0 |
| 3744/580 | 8 | 0.0 |
| 1153 | 8 | 0.0 |
| 1021 | 8 | 0.0 |
| 718/7465/7483 | 8 | 0.0 |
| 779/7045 | 8 | 0.0 |
| 350 | 7 | 0.0 |
| 5477/7845 | 6 | 0.0 |
| 1613 | 6 | 0.0 |
| 10 | 6 | 0.0 |
| 8046/707 | 6 | 0.0 |
| 521 | 5 | 0.0 |
| 509/207 | 5 | 0.0 |
| 2616 | 5 | 0.0 |
| 3904 | 4 | 0.0 |
| 395/8123 | 4 | 0.0 |
| 3142 | 4 | 0.0 |
| 1290/6508 | 3 | 0.0 |
| 2328 | 3 | 0.0 |
| 96 | 3 | 0.0 |
| 816 | 3 | 0.0 |
| 3497 | 3 | 0.0 |
| 5360 | 3 | 0.0 |
| 2096/1760 | 2 | 0.0 |
| 7760 | 2 | 0.0 |
| 913 | 2 | 0.0 |
| 3837 | 2 | 0.0 |
| 1956 | 2 | 0.0 |
| 2990 | 2 | 0.0 |
| 1626 | 2 | 0.0 |
| 3111 | 2 | 0.0 |
| 7314 | 2 | 0.0 |
| 5365/7671 | 2 | 0.0 |
| 1027/5814 | 2 | 0.0 |
| 6063/6064 | 2 | 0.0 |
| 1292/1566 | 2 | 0.0 |
| 6869 | 2 | 0.0 |
| 3675/1094 | 2 | 0.0 |
| 1035 | 2 | 0.0 |
| 574 | 2 | 0.0 |
| 1073 | 2 | 0.0 |
| 1345 | 2 | 0.0 |
| 7047 | 2 | 0.0 |
| 5495/6624 | 2 | 0.0 |
| 5711 | 2 | 0.0 |
| 6610 | 2 | 0.0 |
| 7689/7691 | 2 | 0.0 |
| 1660 | 1 | 0.0 |
| 42 | 1 | 0.0 |
| 8083 | 1 | 0.0 |
| 3691 | 1 | 0.0 |
| 7747 | 1 | 0.0 |
| 3726 | 1 | 0.0 |
| 8048 | 1 | 0.0 |
| 3742 | 1 | 0.0 |
| 1725 | 1 | 0.0 |
| 1558 | 1 | 0.0 |
| 7694 | 1 | 0.0 |
| 136 | 1 | 0.0 |
| 7751 | 1 | 0.0 |
| 3752 | 1 | 0.0 |
| 7847 | 1 | 0.0 |
| 3834 | 1 | 0.0 |
| 8073 | 1 | 0.0 |
| 135 | 1 | 0.0 |
| 699 | 1 | 0.0 |
| 3846 | 1 | 0.0 |
| 3206 | 1 | 0.0 |
| 2098 | 1 | 0.0 |
| 7692 | 1 | 0.0 |
| 4968 | 1 | 0.0 |
| 7745 | 1 | 0.0 |
| 4976 | 1 | 0.0 |
| 7749 | 1 | 0.0 |
| 4981 | 1 | 0.0 |
| 7753 | 1 | 0.0 |
| 4984 | 1 | 0.0 |
| 7840 | 1 | 0.0 |
| 5032 | 1 | 0.0 |
| 7880 | 1 | 0.0 |
| 2099 | 1 | 0.0 |
| 8050 | 1 | 0.0 |
| 5474 | 1 | 0.0 |
| 8077 | 1 | 0.0 |
| 5491 | 1 | 0.0 |
| 2678 | 1 | 0.0 |
| 5494 | 1 | 0.0 |
| 709 | 1 | 0.0 |
| 2106 | 1 | 0.0 |
| 1727 | 1 | 0.0 |
| 5835 | 1 | 0.0 |
| 942 | 1 | 0.0 |
| 356 | 1 | 0.0 |
| 3689 | 1 | 0.0 |
| 6083 | 1 | 0.0 |
| 7693 | 1 | 0.0 |
| 2108 | 1 | 0.0 |
| 7695 | 1 | 0.0 |
| 6612 | 1 | 0.0 |
| 7746 | 1 | 0.0 |
| 6711 | 1 | 0.0 |
| 7748 | 1 | 0.0 |
| 6715 | 1 | 0.0 |
| 7750 | 1 | 0.0 |
| 6868 | 1 | 0.0 |
| 7752 | 1 | 0.0 |
| 2119 | 1 | 0.0 |
| 7754 | 1 | 0.0 |
| 2120 | 1 | 0.0 |
| 7799 | 1 | 0.0 |
| 2233 | 1 | 0.0 |
| 7841 | 1 | 0.0 |
| 6989 | 1 | 0.0 |
| 7849 | 1 | 0.0 |
| 407 | 1 | 0.0 |
| 8043 | 1 | 0.0 |
| 7110 | 1 | 0.0 |
| 8049 | 1 | 0.0 |
| 7258 | 1 | 0.0 |
| 8053 | 1 | 0.0 |
| 7306 | 1 | 0.0 |
| 8075 | 1 | 0.0 |
| 2442 | 1 | 0.0 |
| 8081 | 1 | 0.0 |
| 7471 | 1 | 0.0 |
| 2657 | 1 | 0.0 |
| 7479 | 1 | 0.0 |
| 2767 | 1 | 0.0 |
| 7494 | 1 | 0.0 |
| 2891 | 1 | 0.0 |
| 7497 | 1 | 0.0 |
| 2933 | 1 | 0.0 |
| 7499 | 1 | 0.0 |
| 890 | 1 | 0.0 |
| 7550 | 1 | 0.0 |
| 1766 | 1 | 0.0 |
| 7565 | 1 | 0.0 |
| 3432 | 1 | 0.0 |
| 7592 | 1 | 0.0 |
| 1768 | 1 | 0.0 |
| 7687 | 1 | 0.0 |
| 3646 | 1 | 0.0 |
| 7688 | 1 | 0.0 |
| 7690 | 1 | 0.0 |
| 6979 | 1 | 0.0 |
| 6984 | 1 | 0.0 |
| 3743 | 1 | 0.0 |
| 3747 | 1 | 0.0 |
| Total | 26,677 | 100 |

a. Singletons are unclustered STs of 1 genome each.
