## Supplementary Figures for "Development of a core genome multilocus sequence typing (cgMLST) scheme and life identification number (LIN) code classification system for *Staphylococcus aureus*"

A

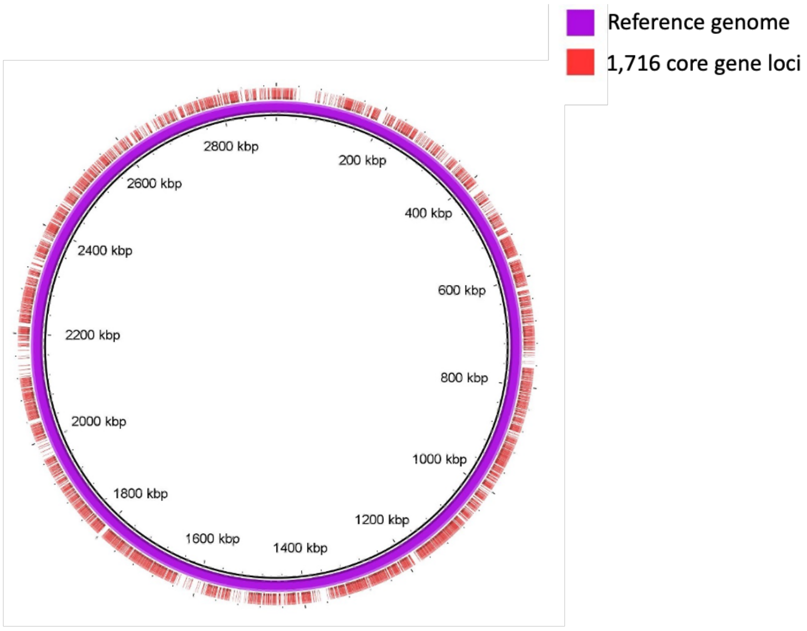


B

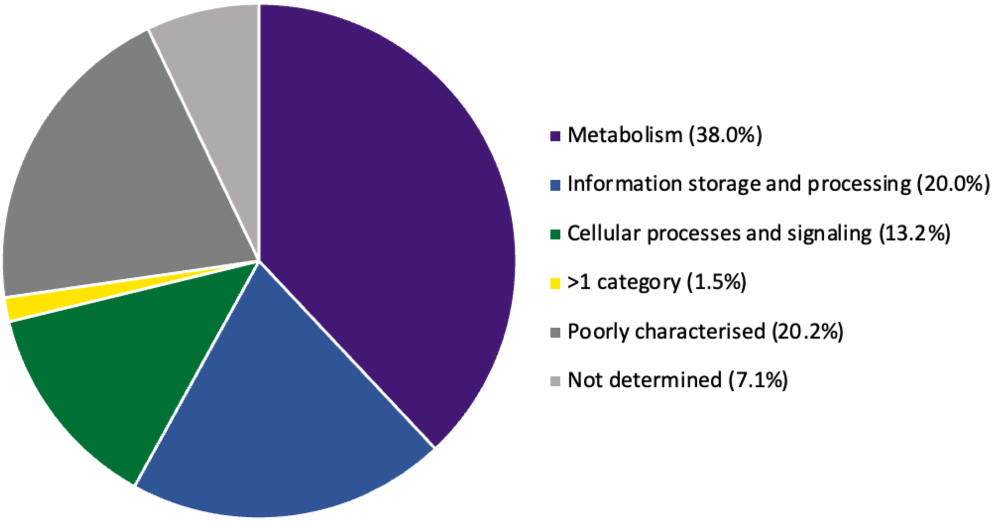


**Supplementary Figure 1. Core gene loci included in the *S. aureus* cgMLST scheme.** A. Distribution of the 1,716 core gene loci across the *S. aureus* reference genome MRSA252 (GenBank accession number BX571856). B. Predicted gene functions of the 1,716 core genes in the cgMLST scheme.

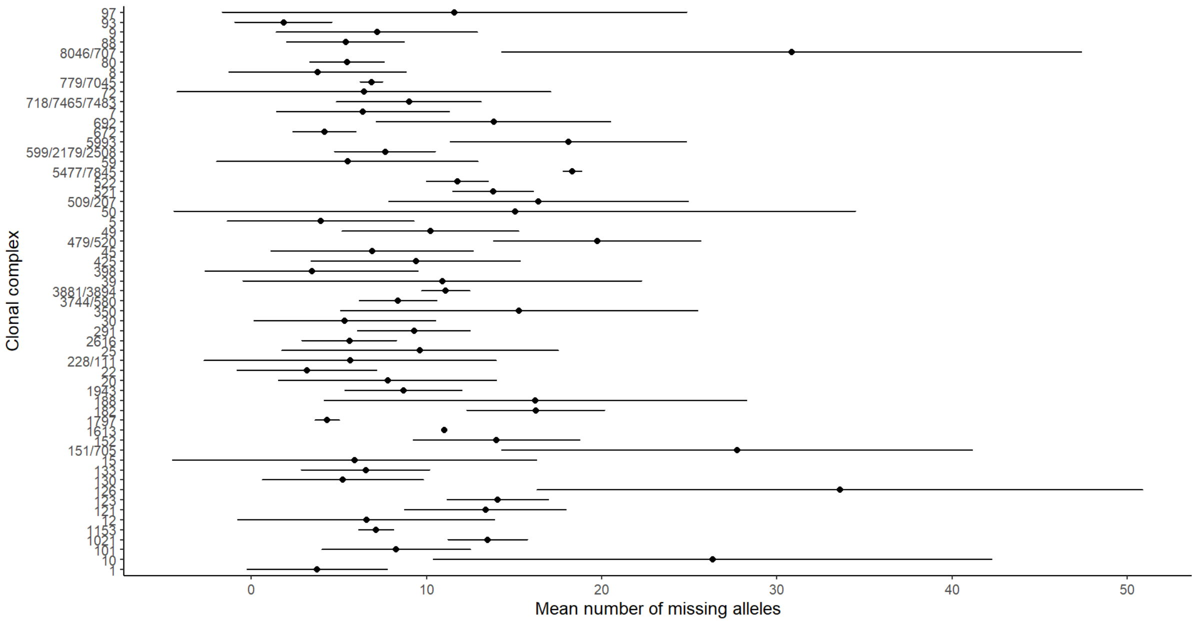


**Supplementary Figure 2. Mean number of missing cgST alleles per genome for each of 57 clonal complexes with 5 or more members.** Horizontal bars represent the standard deviation of the mean.

A

B

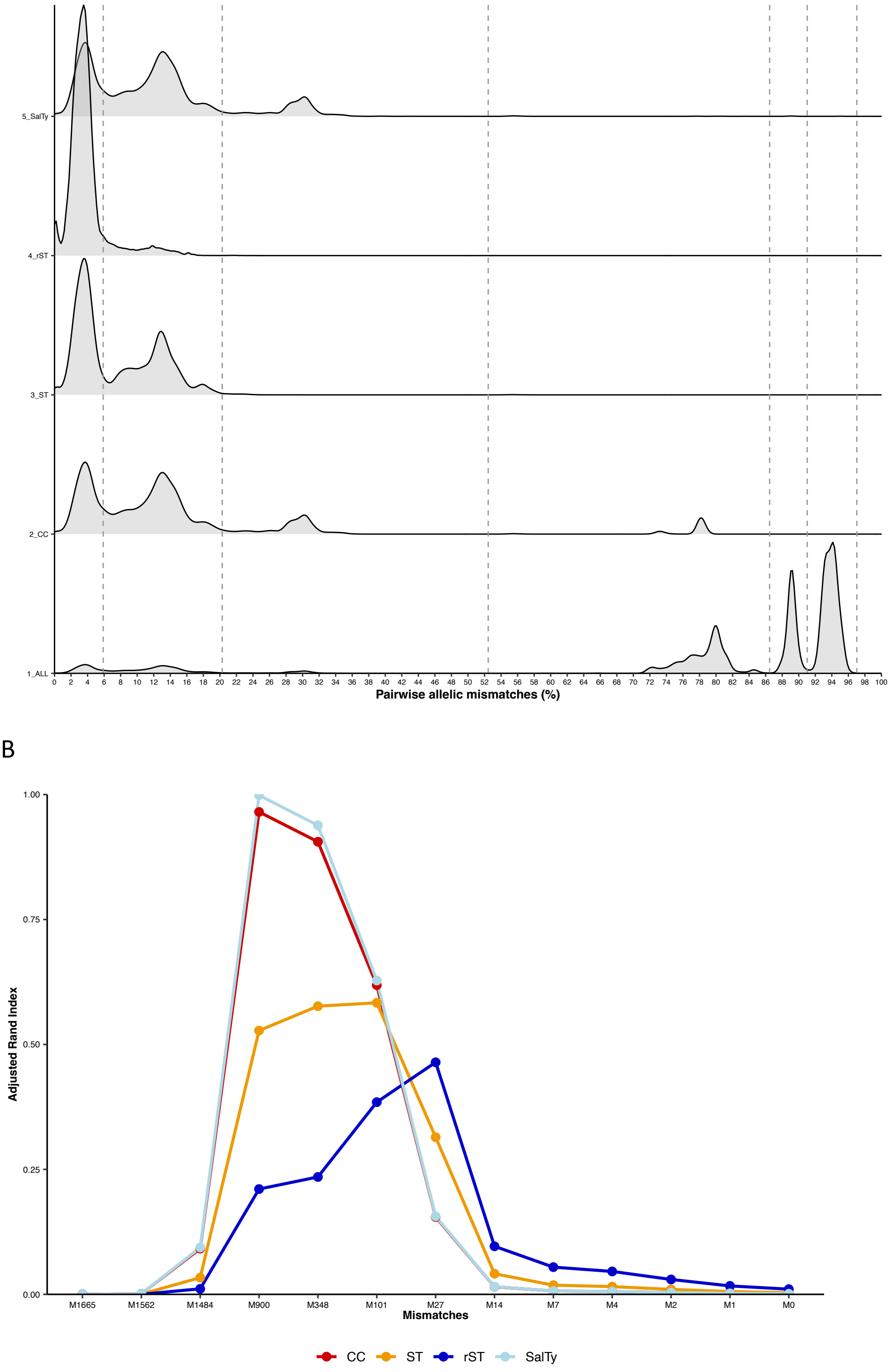


**Supplementary Figure 3. Characterisation of the development dataset of 5,000 *S. aureus* genomes by cgMLST and comparison to other genotyping methods.** Panel A depicts the density distributions of pairwise allelic differences between genomes belonging to a matching rST, ST, CC or SaLTy group. Panel B plots the concordance between clusters of *S aureus* at each of the 13 LIN code thresholds and the corresponding rST, ST, CC or SaLTy designations. (Technical issues prevented a SaLTy analysis of the full genome dataset.)

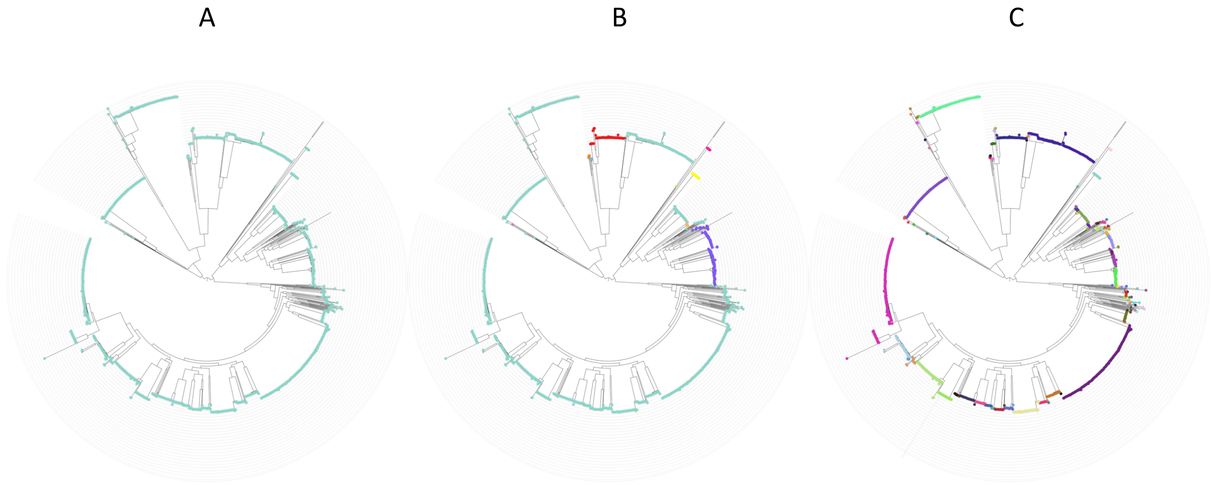


**Supplementary Figure 4. Phylogenetic analysis of *S. aureus* genomes.** The phylogenetic tree was constructed with a concatenated alignment of 1,716 core genome loci from one randomly selected genome representative of each of the 1,558 unique seven-locus sequence types. IQ-TREE was used with the GTR+G model of nucleotide substitution and 1,000 bootstrap replicates. The tree is rooted at the midpoint. The tree was annotated by LIN code groups at three classification levels: (A) LIN code classification threshold 1, (B) LIN code classification threshold 3, and (C) LIN code classification threshold 4 or 'lineage'.
